# Identification of the master sex determining gene in Northern pike (*Esox lucius*) reveals restricted sex chromosome differentiation

**DOI:** 10.1101/549527

**Authors:** Qiaowei Pan, Romain Feron, Ayaka Yano, René Guyomard, Elodie Jouanno, Estelle Vigouroux, Ming Wen, Jean-Mickaël Busnel, Julien Bobe, Jean-Paul Concordet, Hugues Parrinello, Laurent Journot, Christophe Klopp, Jérôme Lluch, Céline Roques, John Postlethwait, Manfred Schartl, Amaury Herpin, Yann Guiguen

**Affiliations:** INRA, UR1037 LPGP, Campus de Beaulieu, Rennes, France.; GABI, INRA, AgroParisTech, Université Paris-Saclay, 78350 Jouy-en-Josas, France.; Fédération d’Ille-et-Vilaine pour la pêche et la protection du milieu aquatique (FDPPMA35), 9 rue Kérautret Botmel – CS 26713 – 35067 RENNES; INSERM U1154, CNRS UMR7196, MNHN, Muséum National d'Histoire Naturelle, 43 rue Cuvier, 75231 Paris Cedex 05, France.; MGX, Univ Montpellier, CNRS, INSERM, Montpellier, France.; Plate-forme bio-informatique Genotoul, Mathématiques et Informatique Appliquées de Toulouse, INRA, Castanet Tolosan, France.; SIGENAE, GenPhySE, Université de Toulouse, INRA, ENVT, Castanet Tolosan, France.; INRA, US 1426, GeT-PlaGe, Genotoul, Castanet-Tolosan, France.; Institute of Neuroscience, University of Oregon, Eugene, OR 97403, USA.; University of Wuerzburg, Physiological Chemistry, Biocenter, 97074 Würzburg, Germany; Comprehensive Cancer Center Mainfranken, University Hospital, 97080 Würzburg, Germany; Hagler Institute for Advanced Study and Department of Biology, Texas A&M University, College Station, Texas 77843, USA.

## Abstract

Teleost fishes, thanks to their rapid evolution of sex determination mechanisms, provide remarkable opportunities to study the formation of sex chromosomes and the mechanisms driving the birth of new master sex determining (MSD) genes. However, the evolutionary interplay between the sex chromosomes and the MSD genes they harbor is rather unexplored. We characterized a male-specific duplicate of the anti-Müllerian hormone (*amh)* as the MSD gene in Northern Pike (*Esox lucius*), using genomic and expression evidences as well as by loss-of-function and gain-of-function experiments. Using RAD-Sequencing from a family panel, we identified Linkage Group (LG) 24 as the sex chromosome and positioned the sex locus in its sub-telomeric region. Furthermore, we demonstrated that this MSD originated from an ancient duplication of the autosomal *amh* gene, which was subsequently translocated to LG24. Using sex-specific pooled genome sequencing and a new male genome sequence assembled using Nanopore long reads, we also characterized the differentiation of the X and Y chromosomes, revealing a small male-specific insertion containing the MSD gene and a limited region with reduced recombination. Our study depicts an unexpected level of limited differentiation within a pair of sex chromosomes harboring an old MSD gene in a wild population of teleost fish, highlights the pivotal role of genes from the *amh* pathway in sex determination, as well as the importance of gene duplication as a mechanism driving the turnover of sex chromosomes in this clade.

**Author Summary:** In stark contrast to mammals and birds, teleosts have predominantly homomorphic sex chromosomes and display a high diversity of sex determining genes. Yet, population level knowledge of both the sex chromosome and the master sex determining gene is only available for the Japanese medaka, a model species. Here we identified and provided functional proofs of an old duplicate of anti-Müllerian hormone (Amh), a member of the Tgf-β family, as the male master sex determining gene in the Northern pike, *Esox lucius*. We found that this duplicate, named *amhby* (Y-chromosome-specific anti-Müllerian hormone paralog b), was translocated to the sub-telomeric region of the new sex chromosome, and now *amhby* shows strong sequence divergence as well as substantial expression pattern differences from its autosomal paralog, *amha*. We assembled a male genome sequence using Nanopore long reads and identified a restricted region of differentiation within the sex chromosome pair in a wild population. Our results provide insight on the conserved players in sex determination pathways, the mechanisms of sex chromosome turnover, and the diversity of levels of differentiation between homomorphic sex chromosomes in teleosts.

## Introduction

The evolution of sex determination (SD) systems and sex chromosomes have sparked the interest of evolutionary biologists for decades. While initial insights on sex chromosome evolution came from detailed studies in *Drosophila* and in mammals [1–5], recent research on other vertebrate groups, such as avian [6,7], non-avian reptiles [8,9], amphibians [10–13], and teleost fishes [14–17], has provided new information that helps us understand the evolution of SD systems and sex chromosomes.

Teleosts display the highest diversity of genetic sex determination systems in vertebrates, including several types of monofactorial and polygenic systems [14,18]. In addition, in some species, genetic factors can interact with environmental factors, most commonly temperature, *i.e.* in *Odontesthes bonariensis* [19], generating intricate sex determination mechanisms. Moreover, sex determination systems in fish can differ between very closely related species, as illustrated in the group of Asian ricefish (genus *Oryzias*) [20– 25], and sometimes even among different populations of one species, like in the Southern platyfish, *Xiphophorus maculatus* [26]. Beside this remarkable dynamic of sex determination systems, the rapid turnover of sex chromosomes in teleosts provides many opportunities to examine sex chromosome pairs at different stages of differentiation. Finally, recent studies on fish sex determination have revealed a dozen new master sex determining (MSD) genes [14,16,27], adding additional insights on the forces driving the turnover of SD systems and the formation of sex chromosomes.

The birth of new MSD genes drives the formation of sex chromosomes and transitions of SD systems. The origin of new MSDs falls into two categories: either gene duplication followed by sub- or neo-functionalization, or allelic diversification [14]. To date, teleosts is the only group where examples of both gene duplication and allelic diversification mechanisms have been found [14–17]. Yet, sex chromosomes with a known MSD gene have only been characterized extensively on the sequence level in two teleost species: the Japanese medaka (*Oryzias latipes*), whose MSD gene originated from gene duplication followed by sub- or neo-functionalization mechanism [28–31], and the Chinese tongue sole, whose MSD gene originated from allelic diversification [32]. Therefore, to form a rich knowledgebase allowing advances of theories of sex chromosomes evolution, additional empirical studies are urgently needed, in particular studies on how the mechanism by which MSDs originate impacts the formation and history of sex chromosomes.

Among identified teleost MSD genes, the *salmonid* MSD gene, named *sdY*, is the most intriguing because it revealed a previously unexpected flexibility in SD pathways in teleosts. While all other currently identified MSDs belong to one of three protein families (SOX, DMRT and TGF-β and its signaling pathway) that were known to be implicated in the SD pathways, the *salmonid* MSD gene *sdY* arose from duplication of an immune-related gene [33]. Despite *sdY* being conserved in the majority of salmonid species [34], it was not found in *Esox lucius*, the most studied member of the salmonid’s sister order the Esociformes [34]. The restriction of *sdY* to the salmonids raised the question of what was the ancestral MSD before the emergence of the “unusual” *sdY.* The first step to answer this question was then to identify the genetic component responsible for sex determination in *E. lucius*.

*E. lucius*, commonly known as the Northern pike, is a large and long-lived keystone predatory teleost species found in freshwater and brackish coastal water systems in Europe, North America, and Asia [35]. It has emerged as an important model species for ecology and conservation because of its pivotal role as a top predator that shapes the structure of local fish communities, and also as a valuable food and sport fish [36]. Consequently, genomic resources have been recently generated for *E. lucius,* including a whole genome assembly anchored on chromosomes [37] and a tissue-specific transcriptome [38]. Yet, little is known about genetic sex determination in *E. lucius* beyond the knowledge that males are the heterogametic sex [39], and its sex locus and MSD gene remain elusive.

In this study, we identified a duplicate of anti-Müllerian hormone (*amh*) with a testis-specific expression pattern as a candidate male MSD gene for *E. lucius*. Using pooled sequencing (pool-seq) reads from a wild population and a new draft genome sequenced with Nanopore long reads, we found limited differentiation between the homomorphic sex chromosomes and that this male-specific duplicate of *amh*, which we call *amhby,* is the located within the Y-specific sequence. Using RAD-sequencing of a family panel, we identified Linkage Group 24 as the sex chromosome and positioned the sex locus in its sub-telomeric region. In addition, we showed that *amhby* has an expression profile characteristic of a male MSD gene and is functionally both sufficient and necessary to trigger testis development, providing robust support for *amhby* as the MSD gene in this species. Finally, through phylogenetic and synteny analyses, we showed that this *amh* duplication took place around 40 million years ago and is lineage-specific and that *amhby* was translocated after its duplication, which likely initiated the formation of the proto-Y chromosome.

Taking advantage of recent advances in functional genomics and sequencing technologies, our study combines the location and characterization of the sex locus and the identification of a master sex determining (MSD) gene with substantial functional validation in a non-model species. Our results expand the knowledge of sex determination genes and provide insight on the evolution of sex chromosomes in teleosts.

## Results

### Identification of a male-specific duplicate of *amh* with testis-specific expression in *E. lucius*

Two *amh* transcripts sharing 78.9% nucleotide identity were identified in the tissue-specific transcriptomes of *E. lucius* ([38], phylofish.sigenae.org), one transcript predominantly expressed in the adult testis with a low expression found also in adult ovary and adult muscle, and the other transcript exclusively expressed in the adult testis (**Fig S1**). PCR amplification on genomic DNA from 221 wild-caught individuals, whose phenotypic sex was determined by gonadal inspection, showed that the genomic sequence of one *amh* copy was present in all phenotypic males and females, while the genomic sequence of the testis-specific *amh* was present in 98% of phenotypic males (157/161) and 0% of phenotypic females (0/60) (**Fig 1A**), demonstrating a highly significant association between this testis-specific copy of *amh* and male phenotype (Chi-squared test, *p*< 2.2e-16), and indicating that the genomic sequence of this testis-specific copy of *amh* transcript is Y chromosome specific. This male-specific *amh* was named *amhby* (Y-chromosome-specific anti-Müllerian hormone paralog b) and the other autosomal gene was named *amha* (*amh* paralog a).

**Figure 1:**
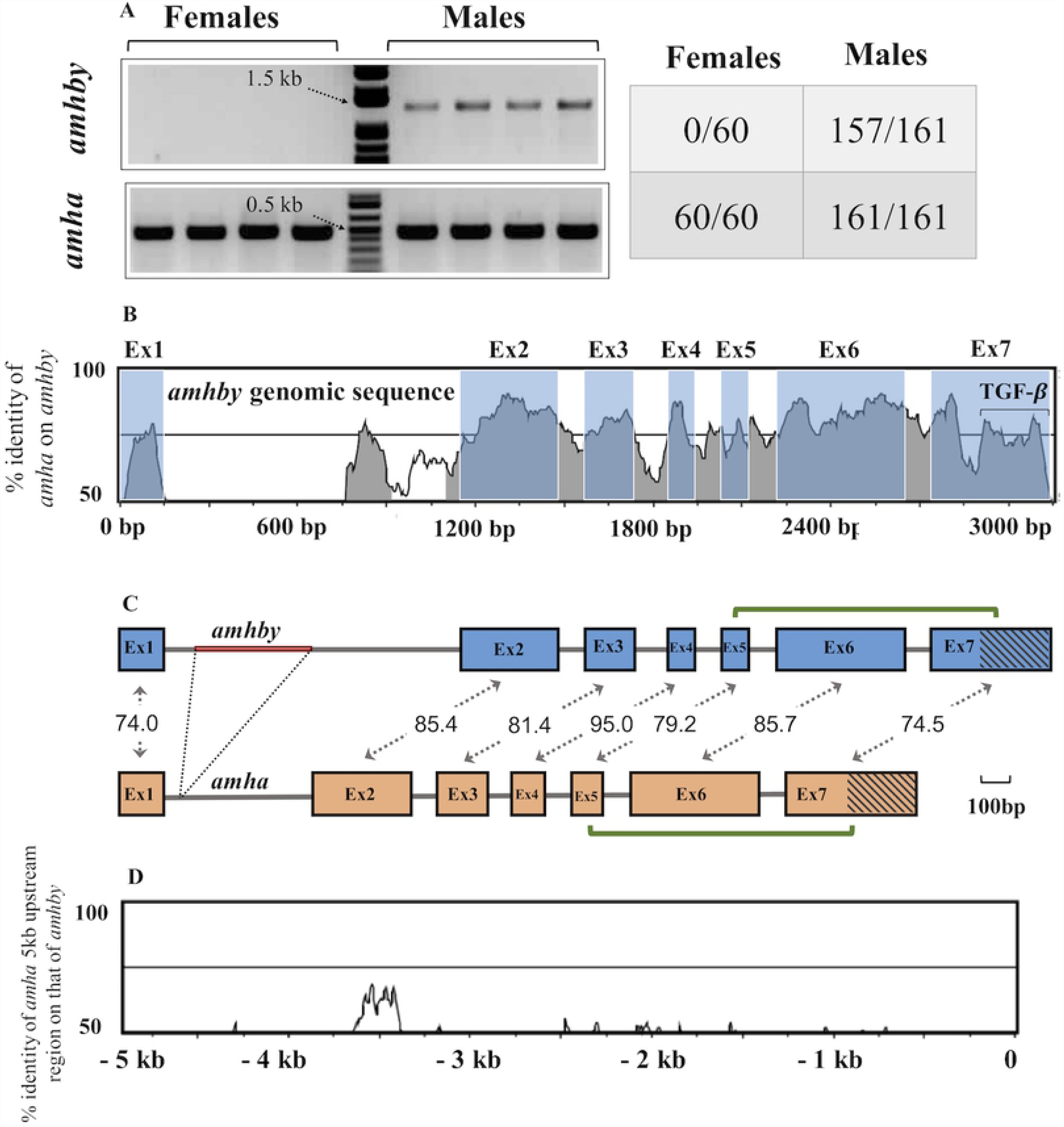
Sequence identity between *amha* and *amhby* and amplification in males and females. A) PCR amplification of *amha* (500 bp band) and *amhby* (1500 bp band) in male (n=4) and female (n=4) genomic DNA samples from *E. lucius*. Each lane corresponds to one individual. The number of tested animals of each sex positive for *amha* and *amhby* is indicated in the table on the right. B) Sequence identify of global pairwise alignment between *amha* and *amhby* genomic sequences with *amhby* sequence as the reference. C) Schematic representation of *amha* and *amhby* gene structure in *E. lucius* from the start to the stop codon. Exon1-Exon7 are represented by green boxes with shared percentage identity indicated and introns are represented by white segment. The red segment in intron 1 represents the *amhby* specific insertion. TGF-β domains are indicated with diagonal lines. D) Global pairwise alignment between 5 kb upstream region of *amha* and *amhby* with *amhby* sequence as the reference. The start codon of each gene is positioned at 0 bp and the number on the y-axis indicates distance from the start codon upstream to the coding sequence.

To compare the genomic regions containing *amha* and *amhby*, clones were isolated from a phenotypic male genomic fosmid library and sequenced. The *amha*-containing fosmid included the entire 5’ intergenic region of *amha* up to the closest gene, *dot1l*, and the *amhby*-containing fosmid included a 22 kb region upstream of *amhby* which contained no coding sequence for other proteins (blastx search against Teleostei, taxid:32443). Nucleotide identity between *amha* and *amhby* exon sequences ranges from 74% to 95% (**Fig 1B and 1C**) and the only gross structural difference between the two genes is a 396 bp specific insertion of a repeated region in *E. lucius* genome in *amhby* intron 1. Little sequence similarity was found between the proximal sequences of the two genes, except for a 1020 bp repetitive element (transposase with conserved domain HTH_Tnp_Tc3_2) (**Fig 1D)**. Predicted proteins contain 580 amino-acids (AA) for *amha* and 560 AA for *amhby*, sharing 68.7% identity and 78.4% similarity. Both proteins have a complete 95 AA C-terminal TGF-β domain with seven canonical cysteines (**Fig S2**), sharing 62.5% identity and 74.0% similarity.

### Assembly of an XY genome containing *amhby* using Nanopore long reads

Blast results revealed that the sequence of the *amhby*-containing fosmid was absent from the available genome assembly (GenBank assembly accession: GCA_000721915.3), which suggests that the sequenced individual could have been a genetic female. Therefore, to characterize the sex locus, we sequenced and assembled the genome of a genetic male with Nanopore long reads. Results from BUSCO show that this new assembly has comparable completeness to that of the reference assembly (**Table S1)**. In this Nanopore assembly, the entire sequence of the *amhby*-containing fosmid was included in a 99 kb long scaffold, tig00003316, from 24,050 bp to 60,989 bp, with *amhby* located from 27,062 bp to 30,224 bp.

### The Y chromosome harbors a small sex locus containing *amhby*

To locate and characterize the sex locus of *E. lucius*, we first aligned sex-specific pool-seq reads from 30 phenotypic males with *amhby* and 30 phenotypic females without *amhby* on the available Northern pike reference assembly (GenBank assembly accession: GCA_000721915.3), which we suspected to be an XX genome, and computed the number of male-specific SNPs (MSS) in a 50 kb non-overlapping window across the genome (**Fig 2A**). Genome average was 0.08 MSS per window, and two windows located on unplaced scaffold 1067 contained 136 and 89 MSS respectively, close to four times the number of MSS in the next highest window (23 MSS). A further analysis with a higher resolution (i.e., 2.5 kb windows) revealed that the majority (94%) of the MSS on scaffold 1067 is restricted to a 40 kb region located between ~ 143 kb and ~ 184 kb on this scaffold. Together, these results indicate that the sex locus is located on scaffold 1067, and Y-derived sequences show strong differentiation from the XX genome only in a small 40 kb region on this scaffold (**Fig 2B, Fig S3**).

**Figure 2:**
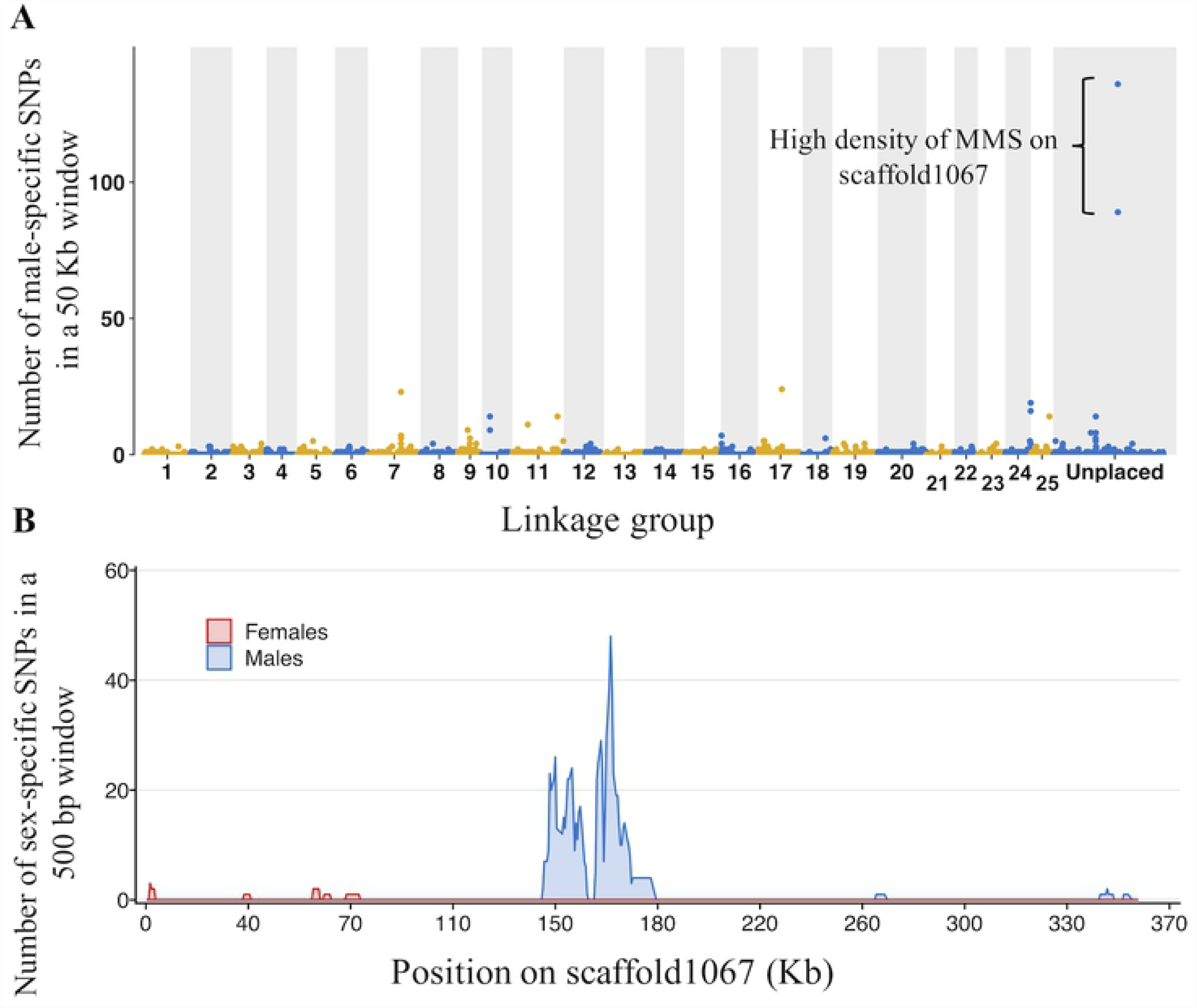
Regions enriched with male-specific SNPs (MMS) in the genome of *E. lucius* based on pooled sequencing analysis. **A**: Number of MSS in a 50 kb non-overlapping window is plotted for each linkage group (LG) and all unplaced scaffolds in the reference genome (GenBank assembly accession: GCA_000721915.3). The two highest peaks of MMS density are located on unplaced scaffold1067. **B**: Number of male and female-specific SNPs in 50 kb non-overlapping windows is plotted along scaffold1067. The male data are represented in blue and female in red. The ~ 40 kb region located between ~ 143 kb to ~ 184 kb (11% of the scaffold) contained 211 MMS, showing the strongest differentiation between males and females.

In a second step, to identify male-specific regions, we aligned the pool-seq reads on the XY Nanopore assembly and identified 94 non-overlapping 1 kb regions covered only by reads from the male pool (MR1k). These regions were located on only three Nanopore contigs (tig00003316 = 53 MR1k, tig00003988 = 22 MR1k, tig00009868 = 19 MR1k). In contrast, there were only four non-overlapping 1 kb regions covered only by reads from the female pool, each located on a different contig, indicating a low false discovery rate for sex-specific regions with our method. Moreover, we found no MR1k from the same analysis on the reference assembly (GenBank assembly accession: GCA_000721915.3), further confirming that the individual sequenced for this assembly has an XX genotype. Blasting results on the three MR1k-containing Nanopore contigs showed that *amhby* is the only protein coding gene apart from transposable elements (TEs) associated proteins in the sex locus of *E. lucius* (**Table S2**).

To better delimitate the sex locus, we searched for Y-specific sequences, defined as regions with no or little female reads mapped and male reads mapped at a depth close to half of the genome average, on the three MR1k-containing contigs from the Nanopore assembly (**Supplementary File 1**, **Fig S4**). In total, we identified ~ 180 kb of Y-specific sequences.

Taken together, these results indicate that the sex locus of *E. lucius,* is ~ 220 kb in size including ~ 180 kb of Y-specific sequence and ~ 40 kb of X/Y differentiated region, and that *amhby* is the only non-TE, protein-coding gene in this locus.

### The sex locus of *E. lucius* is located in the sub-telomeric region of LG24

The pool-seq results located the sex locus on unplaced scaffold 1067. To identify the sex chromosome in the genome of *E. lucius*, we generated RAD-Seq data from a single full-sibling family of *E. lucius* with two parents, 37 phenotypic male offspring, and 41 phenotypic female offspring. In total, 6,512 polymorphic markers were aligned to the 25 nearly chromosome-length linkage groups and 741 polymorphic markers were aligned to unplaced scaffolds of the reference assembly of *E. lucius* (GenBank assembly accession: GCA_000721915.3). Genome-wide average F_st_ between males and females was 0.0033. Only LG2 (F_st_ = 0.006) and LG24 (F_st_ = 0.025) had a higher average F_st_ between males and females than the genome average, and only markers on LG24 showed genome-wide significant association with sex phenotype (**Fig 3A**), indicating that LG24 is the sex chromosome of *E. lucius*. The F_st_ of markers aligned to LG24 gradually increased along the length of the chromosome and reached 90 times the genome average F_st_ (0.38) at the distal end of the chromosome (**Fig 3B)**, and the strongest association with sex was also identified for markers aligned to this region (**Fig 3C)**, pinpointing the location of the sex locus to the sub-telomeric region of LG24. In addition, with our parameters, 32 non-polymorphic RAD markers were found only in the father and all male offspring, 15 of which could be aligned to the reference XX genome (**Table S3**): five (33%) to a region located distal to 22.1 Mb on LG24; one (7%) at position 17.3 Mb on LG24; seven (22%) to unplaced scaffold 0213, and two (6%) to other LGs. Moreover, two of the other 17 male-specific markers which could not be aligned to the reference genome aligned perfectly to the sequence of *amhby*, showing that *amhby* has a strict father-to-son inheritance pattern. Collectively, these results locate the sex locus, containing *amhby*, in the sub-telomeric region of LG24 of *E. lucius*. In addition, unplaced scaffold 0213, which is enriched in male-specific RAD markers, and unplaced scaffold 1067, which is enriched with male-specific SNPs identified with the pool-seq analysis, should also be placed in the sub-telomeric region of LG24.

**Figure 3:**
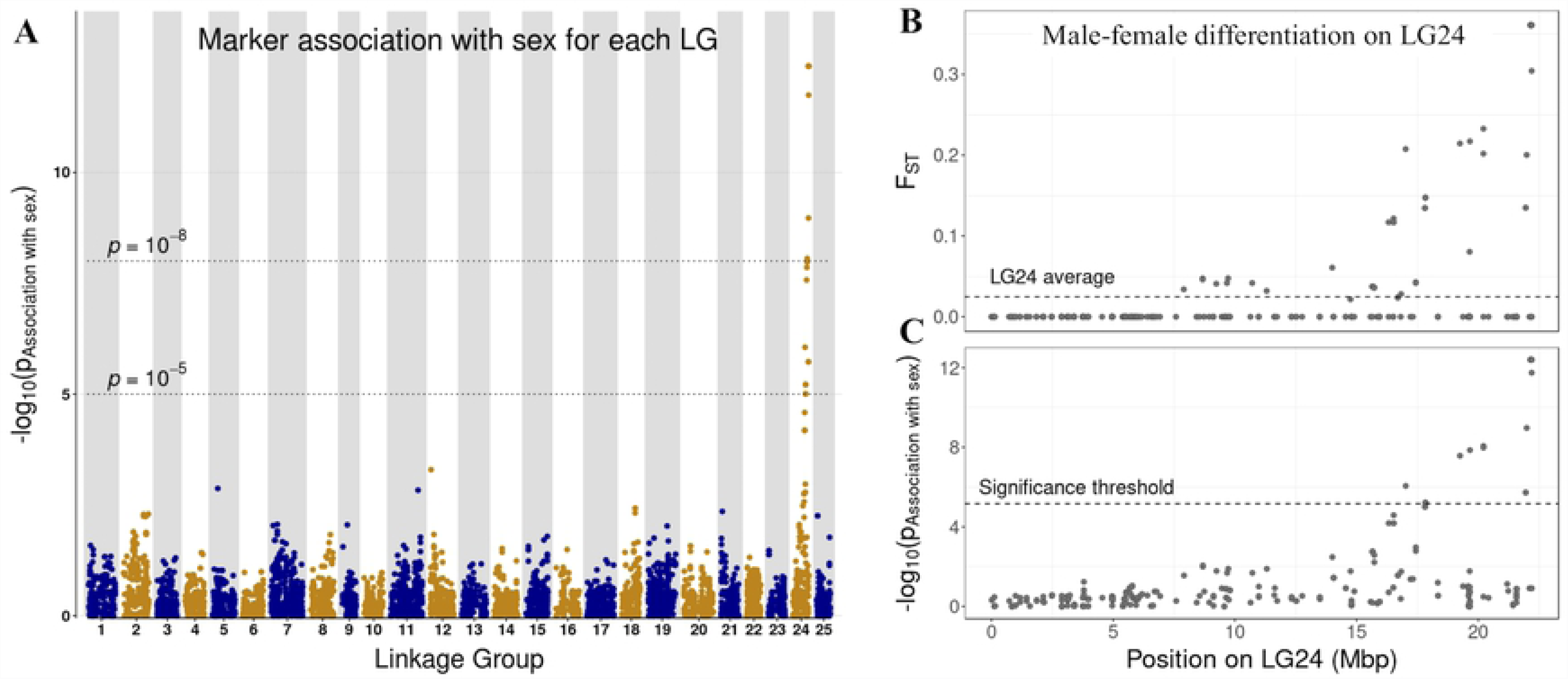
A: Rad-Seq marker association with sex across the 25 linkage groups of *E. lucius* genome. The log_10_ of p-value from the association test between each marker and sex phenotype is mapped against the linkage group (LG) they belong. Linkage group 24 showed a concentration of markers significantly associated with sex phenotype. The dotted horizontal lines are indications for genome-wide significance level from the association test, with the lower dotted line indicating p-value of 1*10^-5^ and higher dotted line indicating p-value of 1*10^-8^. **B**: Differentiation of polymorphic markers demonstrated by F_ST_ between males and females and their association with sex phenotypes on LG24. Between male and female F_ST_ value is plotted against the mapped position of each marker on the upper panel. Average F_ST_ value of markers on LG24 is presented as the dotted line (higher panel). **C**: The log10 of p-value from the association test of each marker with sex phenotype is mapped against the mapped position on the lower panel. Significant threshold of p-value for this test is presented as the dotted line (lower panel). Both between-sex F_ST_ and marker association with sex indicate that the sex locus is near the end of LG24.

### *amhby* is expressed prior to molecular gonadal differentiation in male *E. lucius*

To characterize the temporal and spatial expression of *amhby* in relation to the molecular and morphological differentiation between male and female gonads, both quantitative PCR (qPCR) and *in-situ* hybridization (ISH) were performed.

Expression of *amhby*, *amha,* three other genes (*drmt1*, *cyp19a1a*, and *gsdf*) known for their role in gonadal sex differentiation, and *amhrII*, the putative receptor for the canonical *amh,* was measured by qPCR at four time points from 54 days post-fertilization (dpf) to 125 dpf, prior to the onset of gametogenesis. The entire trunks were used for the first three time points when the gonads were too small to be isolated, and only gonads were used at 125 dpf for both males and females. Expression of *amha* was detected in both males and females starting from 75 days post-fertilization (dpf), with a significantly higher expression in males than in females at 100 dpf (Wilcoxon signed-rank test, *p*=0.043) (**Fig 4A**). In contrast, expression of *amhby* was detected only in males starting from 54 dpf and increasing exponentially thereafter till 125 dpf (R²=0.79) (**Fig 4B**). Expression of *drmt1*, *cyp19a1a*, and *gsdf* was only detected from 100 dpf onwards, *i.e.* much later than the first detected expression of *amhby* (**Fig S5**). Moreover, among these three genes, only *cyp19a1a* showed significantly different expression between sexes with a higher expression in females at 100 dpf (Wilcoxon signed-rank test, *p*=0.014). Expression of *amhrII* was not detected until 100 dpf and did not differ significantly between sexes at any stage **(Fig S5)**.

**Figure 4:**
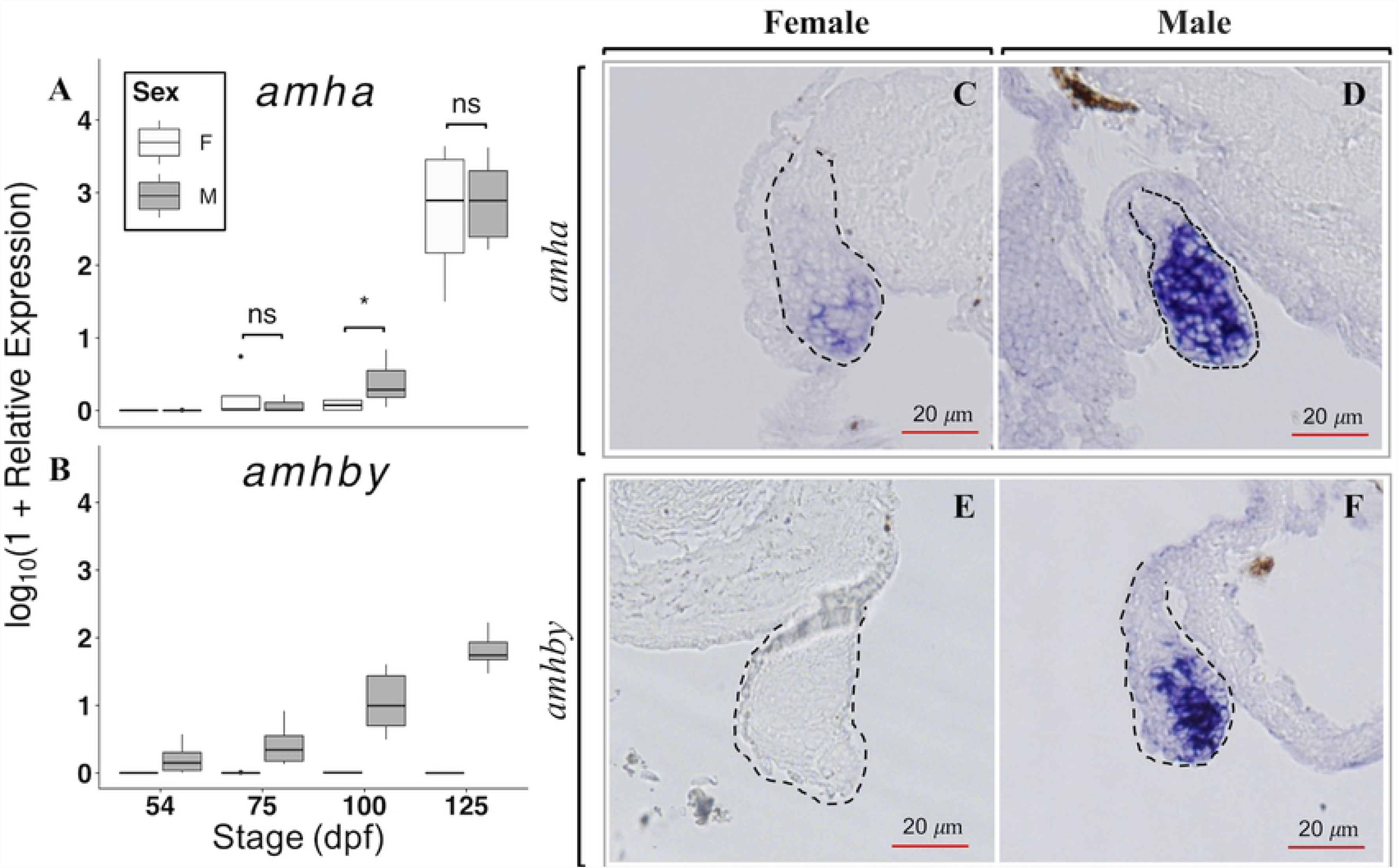
Temporal and spatial expression of *amha* and *amhby* mRNA in male and female developing gonads. **A-B**: Boxplots showing the first quantile, median, and the third quantile of the temporal expression of *amha* (A) and *amhby* (B) during early development of *E. lucius* measured by qPCR. Outliers are displayed as a dot. The mRNA expression of *amha* and *amhby* were measured at 54, 75, 100, and 125 days post fertilization (dpf) in male and female samples of *E. lucius* and the log_10_ of their relative expression is presented on the graph. Significant P-values (<0.05) for Wilcoxon signed rank test between male and female expression at each time point are indicated by * and ‘ns’ indicates ‘non-significant P-values. Statistical tests were not performed on *amhby* expression between sexes because of the complete absence of *amhby* expression in females. **C-F**: *In situ* hybridization on histological sections revealed the localization of *amha* in both 80 dpf female (C) and male (D) gonads, with a stronger expression in male gonads. A high *amhby* mRNA expression is detected in the male gonad (F), with no signal detected in female gonad (E). The red scale bars denote 20 *𝜇*m and the dashed lines outlines the gonadal sections.

Expression of *amha* and *amhby* was also characterized by *in-situ* hybridization (ISH) performed on histological sections of the entire trunk of male and female *E. lucius* sampled at 80 dpf. Expression of *amha* was detected in the gonads of both female (**Fig 4C**) and male (**Fig 4D**) samples, but the signal was much stronger in male gonads. In contrast, expression of *amhby* was strong in male gonads **(Fig 4F)** but not detected in female gonads **(Fig 4E)**, confirming the specificity of the probe for *amhby*. In addition, no morphological differences were observed between male and female gonads at 80 dpf, even though expression of *amhby* was already detected by qPCR and by ISH at this stage. Our ISH results show that the expression of both *amha* and *amhby* is high in male gonads before the first signs of histological differentiation between male and female gonads.

Collectively, these results show that *amhby* is expressed in the male gonads prior to both molecular and morphological sexual-dimorphic differentiation of gonads in *E. lucius*.

### The *amhby* gene is both necessary and sufficient to trigger testicular differentiation

To further investigate the functional role of *amhby* in initiating testicular development, we performed both loss-of-function and gain-of-function experiments.

We knocked out *amhby* using three pairs of TALENs targeting exon 1 and exon 2 of *amhby* (**Fig S4A**). Only the T2 TALEN pair targeting exon 1 was effective in inducing deletions in the *amhby* sequence. Overall, 12 of 36 (33.3%) surviving G0 males possessed a disrupted *amhby* that resulted in truncated proteins (**Fig S4B)**. G1 XY offspring obtained from three *amhby* mosaic G0 males crossed with wild-type females were maintained until the beginning of testicular gametogenesis at 153 dpf and then processed for histology. Gonads from 23 G1 *amhby* positive XY mutants were compared with those of wild-type control XY males (N=4) and control XX females (N=4) of the same age. Control animals developed normal ovary and testis (**Fig 5A and 5B)**, but all 23 XY F1 *amhby* mutants failed to develop a normal testis. Among these 23 G1 XY mutants, 20 (87%) showed complete gonadal sex reversal, characterized by the formation of an ovarian cavity and the appearance of previtellogenetic oocytes (**Fig 5C)**; the three (13%) remaining mutants developed potentially sterile gonads with no clear ovarian nor testicular structure.

**Figure 5:**
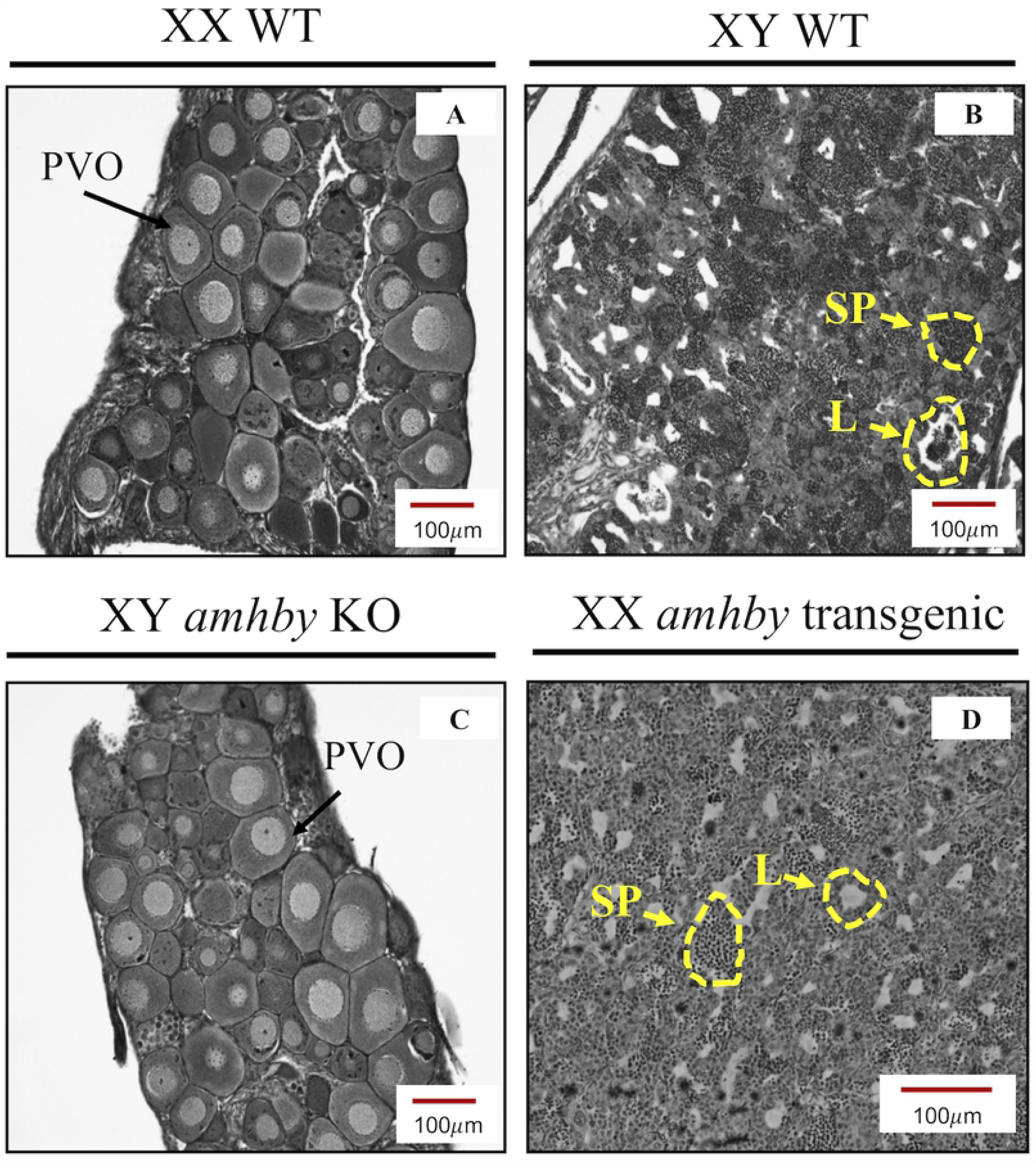
Gonadal phenotypes of *E. lucius* in *amhby* knockout (KO) and additive transgenesis experiments. Gonadal histology of a representative control XX female (A), control XY male (B), an *amhby* KO XY male (C), and an *amhby* transgenic XX mutant female (D). The *amhby* KO XY male (C) developed ovaries with oocytes and an ovary cavity, indistinguishable from the ovary of the control females (A). The *amhby* transgenic XX mutant female (D) developed testis with clusters of spermatozoids and testicular lobules identical to that of the control males (B). **PVO**: previtellogenic oocytes; **SP**: spermatozoids; **L**: testicular lobules; **T**: testis; and **O**: ovary.

To investigate whether *amhby* alone is sufficient to trigger testicular development, we overexpressed *amhby* in XX genetic females. Two G0 XY mosaic transgenic males possessing the *amhby* fosmid were crossed to wild-type females, and 10 G1 XX offspring carrying the *amhby* fosmid were maintained along with control wild-type siblings until the beginning of testicular gametogenesis at 155 dpf. Upon histological analysis of the gonads, all ten (100%) XX transgenics carrying *amhby* fosmid developed testis with testicular lumen and clusters of spermatozoids (**Fig 5D)**, while all 12 control genetic males developed testis and 19 of 24 (79%) control genetic females developed ovaries. The other five control genetic females (21%) developed testes. Such sex reversal was also observed in the natural population at a rate of 2%, and this might have been exacerbated by culture conditions, a phenomenon previously documented in other teleosts [40]. Despite this effect, the XX transgenics with *amhby* fosmid had a significantly higher rate of sex reversal than their control female siblings raised in identical condition (Chi-squared test, *p*= 0.0001148).

Taken together, these results show that *amhby* is both necessary and sufficient to trigger testicular development in *E. lucius*, and further support the functional role of *amhby* as the MSD gene in this species.

### A chromosomal translocation involved *amhby* after a lineage-specific duplication of *amh*

To determine the origin of the two *E. lucius amh* paralogs, we generated a map of conserved syntenies for *amh* in several teleost species, including the spotted gar (*Lepisosteus oculatus*) as outgroup (**Fig 6A**). Genes located upstream (*i.e. dot1l, ell,* and *fkbp8*) and downstream (*i.e. oazla*) of *amha* on *E. lucius* LG08 showed conserved synteny in all teleost species included in the analysis, indicating that LG08 is the conserved location of the *E. lucius* ancestral *amh*, now called *amha*, and that *amhby* evolved from a duplication of *amha* that was later translocated to the sub-telomeric region of the future sex chromosome, LG24. We estimated the duplication event to be ~ 38 and ~50 million years old (**Supplementary file 1**), and found no homology between the ~ 180 Kb Y-specific sequence identified in the Nanopore assembly and the sequence of LG08 from the reference assembly, besides the two *amh* genes, suggesting that this translocation event is also likely to be ancient.

**Figure 6:**
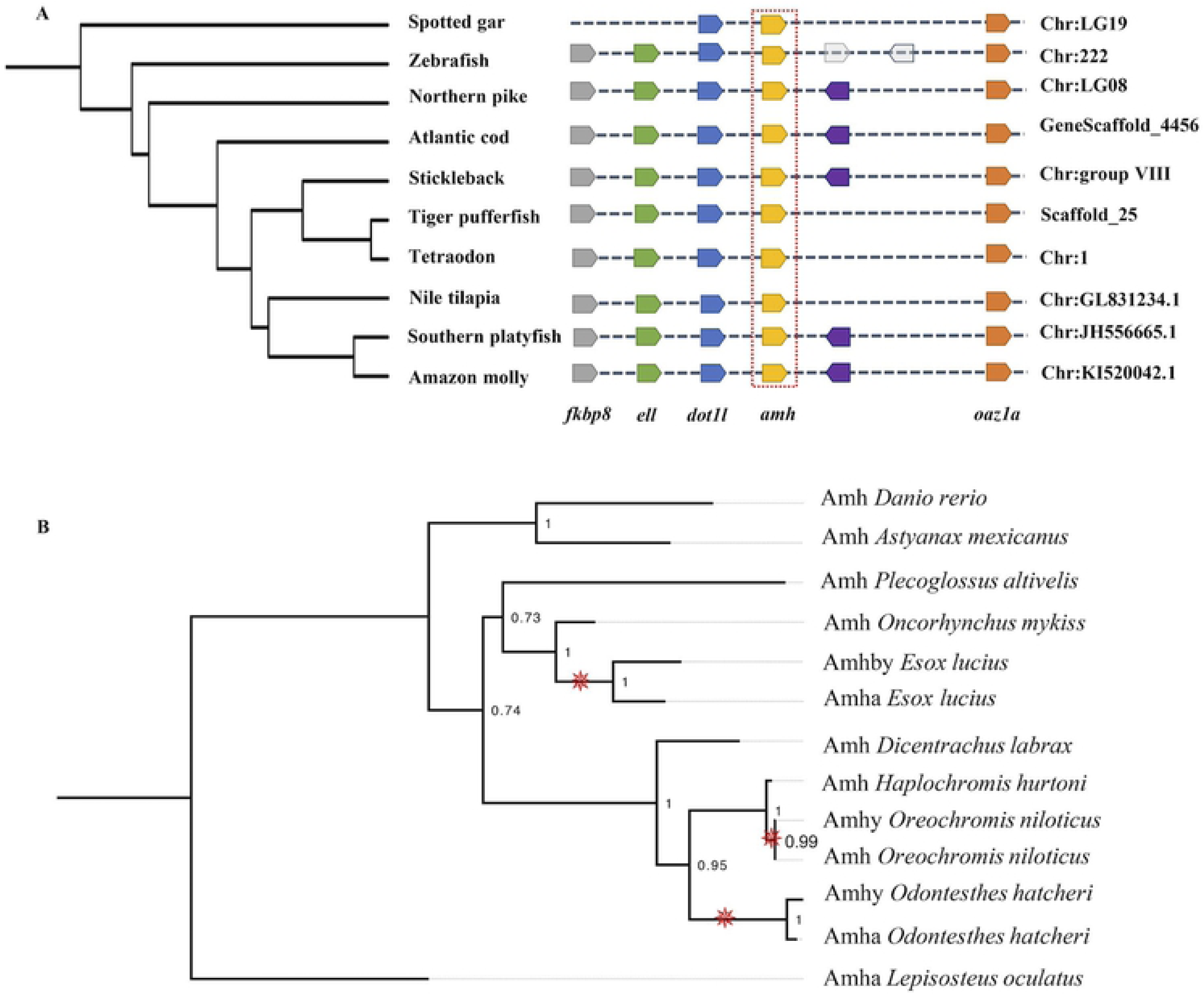
Evolution of Amh in teleosts. **A**: Synteny map of genomic regions around *amh* genes (highlighted by the red box) in teleosts. Orthologs of each gene are shown in the same color and the direction of the arrow indicates the gene orientation. Ortholog names are listed below and the genomic location of the orthologs are listed on the right side. For *Esox lucius*, *amha*, which is located on LG08, is used in this analysis. **B**: Phylogenetic reconstruction of teleost Amh protein orthologs. The phylogenetic tree was constructed with the maximum likelihood method (bootstrap=1000). Numbers at tree nodes are bootstrap values. Spotted gar (*Lepisosteus oculatus*) Amh was used as an outgroup. Branches with Amh duplication are indicated by the red pinwheels.

Prior to the discovery of *amhby* in *E. lucius*, male-specific duplications of *amh* have been identified in Patagonian pejerrey (*Odontesthes hatcheri*) [41] and in Nile tilapia (*Oreochromis niloticus*) [42]. To test whether these duplications have a shared origin, we constructed a phylogeny of Amh from nine teleost species, including these three species with male-specific *amh* duplications, and including spotted gar Amh as outgroup (**Fig 6B**). In this protein phylogeny, each sex-specific Amh paralog clusters as a sister clade to its own species’ ‘canonical’ Amh with significant bootstrap values, indicating that these three pairs of *amh* paralogs were derived from three independent and lineage-specific duplication events.

## Discussion

### *amhby*, the male-specific duplicate of *amh*, is the master sex determination gene in *E. lucius*

Since the discovery of *dmrt1bY* in the Japanese rice fish [28,29], the first identified teleost master sex determination gene, studies in teleosts have unveiled a dozen novel genes as master regulators for sex determination [14–16,27]. Interestingly, many of these master sex determining genes belong to the TGF*-β* superfamily. To date, this finding has been mostly restricted to teleosts, highlighting the crucial role of TGF*-β* signaling in the sex determination pathway in this vertebrate group. In the present study, we identified an old duplicate of *amh*, a member of the TGF*-β* superfamily, as the male MSD gene in *E. lucius*. Results from genotyping demonstrate a strong and significant association of *amhby* with male phenotype. RNA-seq, qPCR and ISH showed that *amhby* is expressed in the male gonadal primordium before histological testis differentiation, thus fulfilling another criterion for being an MSD. Furthermore, knockout of *amhby* leads to complete gonadal sex reversal of XY mutants, while overexpression of *amhby* in XX animals leads to the development of testis, demonstrating that *amhby* is both necessary and sufficient for testicular differentiation. Together, these independent lines of evidence provide strong confidence that *amhby* is the MSD in *E. lucius*.

This work provides a third functionally validated case of an *amh* duplicate evolving into the MSD gene in a teleost species, along with the Patagonian pejerrey, *Odontesthes hatcheri*, [41], and the Nile tilapia, *Oreochromis niloticus*, [42]. Besides these three examples, association of *amh* duplicates with phenotypic sex was also found in other teleosts: in *O. bonariensis*, the sister species of the Patagonian pejerrey, a male-specific *amhy* was found to interact with temperature in determining sex [19], and in the ling cod, *Ophiodon elongatus,* a male-specific duplicate of *amh* was also identified using molecular marker sequences [43]. More recently, a duplicated copy of *amh* was found in an *Atheriniformes* species, *Hypoatherina tsurugae,* and is suspected to be involved in male sex determination [44]. Our phylogenetic analysis on Amh sequences from several teleost species revealed that the three confirmed male-specific Amh duplications are independent, lineage-specific events rather than the product of shared ancestry. This finding supports the “limited option” hypothesis for master sex determining genes [45] and makes *amh* duplicates the most frequently and independently recruited master sex determining genes identified in any animal group so far.

Among the three teleost species with confirmed *amh* duplicate MSD genes, *E. lucius* has the highest degree of divergence in sequence between paralogs, with ~ 79.6% genomic sequence identity on average. In the Nile tilapia, *amhy* is almost identical to the autosomal *amh*, differing by only one SNP [42]. In the Patagonian pejerrey, the shared identity between paralogs ranges from 89.1% to 100% depending on the exon [41]. Because of the low divergence between the two paralogous sequences in the Patagonian pejerrey and the Nile tilapia, the new MSD function of *amhy* was, in both cases, mainly attributed to its novel expression pattern. Yet, low sequence divergence, as little as one amino acid-changing SNP in *amhrII* alleles in *Takifugu*, was shown to be sufficient to impact the signal transduction function of a protein [46]. In *E. lucius*, *amhby* also have an expression profile different from that of *amha*, likely due to a completely different sequence in the putative promoter region. However, because of the relatively high level of divergence between *amha* and *amhby* sequences in *E. lucius*, especially in the C-terminal bioactive domain of the proteins [47], it is tempting to hypothesize that the two proteins could have also diverged in their function, with for instance a different affinity for their canonical AmhrII receptor or even the ability to bind to different receptors, leading to divergences in their downstream signaling pathways. Because we estimated the duplication of *amh in E. lucius* to be between ~ 38 and ~ 50 million-year-old, the long divergence time between the two paralogous *amh* genes potentially provided opportunities for the accumulation of these sequence differences. Further functional studies would be required to unravel the downstream signaling pathways of Amha and Amhby in *E. lucius* to better understand the mechanisms leading to the novel function of *amhby* as the MSD gene in this species.

Surprisingly, the sequence of *amhby* and the ~ 90 kb male specific sequence flanking *amhby* are absent from the reference genome assembly sequenced from DNA of a phenotypically male sample of *E. lucius* [37], suggesting that this sequenced individual had an XX genotype. The absence of male specific regions when mapping pool-seq reads to this assembly further confirms that the reference genome sequence likely comes from an XX genetic individual. This apparent discordance of genotype and phenotype could be due the difficulty in correctly identifying the phenotypic sex of an individual with immature gonads, as Northern pikes do not display sexual dimorphic external traits. Alternatively, the sequenced individual may have been a sex-reversed genetic female, as found in a low proportion (~ 2%) in a wild population and at higher proportion (~ 20%) in captivity raised Northern pikes.

### The birth of the MSD and sex chromosomes in *E. lucius*

The analysis of the genomic neighborhood of both *amh* duplicates showed that *amha* is located on LG08 in a cluster of genes regulating sexual development and cell cycling with conserved synteny in teleosts [48,49], while *amhby* is located near the telomeric region of LG24 with no other identified gene besides transposable elements within at least 99 kb in its close vicinity. These results indicate that *amha* is likely to be the ancestral *amh* copy in *E. lucius*, and that *amhby* was translocated near the telomere of the ancestor of LG24 after its duplication. This scenario fits the description of a proposed mechanism of sex chromosome emergence and turnover through gene duplication, translocation and neofunctionalization [31,50]. Following this model, the translocation of a single copy of *amh* into another autosome triggered the formation of proto sex chromosomes, possibly because the newly translocated genomic segment containing the *amh* copy halted recombination with the X chromosome *ab initio* due to a complete lack of homology. This mechanism came to light after the discovery of *dmrt1bY* in the Japanese medaka, which acquired a pre-existing *cis*-regulatory element in its promoter through a transposable element [51]. Besides *dmrt1bY*, the only other well-described case of sex chromosome turnover via gene duplication, translocation and neofunctionalization is the salmonid *sdY* gene, which maps to different linkage groups in different salmonid species. Our study provides a third empirical example of gene duplication and translocation giving rise to new MSD genes and bolster the importance of this mechanism in the birth and turnover of sex chromosomes.

Theories of sex chromosomes evolution predict suppression of recombination around the sex determining locus, eventually leading to degeneration of the locus because it lacks the ability to purge deleterious mutations and repeated elements [52,53]. Here, we found that only a small part (around 220 kb) of the sex chromosome shows differentiation between X and Y chromosome, suggesting that this sex locus encompasses less than 1% of LG24 in *E. lucius*. Sex chromosome differentiation has been characterized in a few other teleost species, including Chinese tongue sole [32], Japanese medaka [30,54], stickleback [55], Trinidad guppy [56] and a few cichlid species [57,58]. Compared to these examples, we found that the Northern pike displays a very limited region of suppressed recombination between the sex chromosomes. A usual explanation for a small sex locus is that the rapid transition of sex chromosomes frequently observed in teleosts, facilitated by a duplication and translocation mechanism, can readily produce neo-sex chromosomes showing little differentiation except for the recently acquired MSD genes, as was demonstrated in the salmonids with the vagabond MSD gene *Sdy* [34,59,60]. However, the ~ 40 million year of divergence time between *amhby* and *amha*, and the lack of homology between the sequence from the sex locus and that of the vicinity of *amha* on LG08, suggest that the sex locus of *E. lucius* is not nascent. Further comparative studies that include sister species in the same clade will be needed to better estimate the age of the MSD gene and the sex chromosome, but a nascent sex locus is likely not the explanation for the restricted region suppression of recombination between the X and Y chromosome of *E. lucius*. On the other hand, old yet homomorphic sex chromosomes have been observed, for instance in ratite birds [61] and in pythons [62]. Furthermore, in *Takifugu*, a single SNP conserved for 30 million years determines sex and the rest of the sex chromosomes does not show evidence of suppressed recombination, raising the possibility that decay is not the only possible fate for sex chromosomes [46]. One mechanism for the maintenance of a small sex locus has been proposed in the Japanese rice fish, where long repeats flanking the sex locus on the Y chromosome may recombine with the same repeats on the X chromosomes, thus hindering the spread of suppression of recombination around the MSD gene [50]. We found a slight enrichment of repeated elements on the sequences from the sex locus of *E. lucius,* however a better assembly of the sex locus would be needed to investigate whether a similar mechanism could have contributed to the very limited differentiation between X and Y chromosomes in *E. lucius*.

To date, empirical support for non-decaying sex chromosome is still rare, possibly due to the difficulties in identifying small differences between sex chromosomes, and because environmental factors affecting the sex determination pathways could weaken the association between genotype and phenotype, particularly in non-endothermic vertebrates. However, our study, as well as other recent effort in characterizing sex loci in non-model species [63], shows that using current population genomic approaches, we can now relatively easily identify and characterize small sex loci. Future studies surveying sex chromosomes at various stages of differentiation and understanding the factors influencing their level of differentiation will form a new empirical basis to update the current models of sex chromosome evolution.

In conclusion, our study identified an old duplication of *amh* in *E. lucius* which generated a Y-chromosome-specific copy that we named *amhby*. We showed that *amhby* is functionally necessary and sufficient to trigger testicular development, and is expressed in the male gonadal primordium, fulfilling key requirements for a classic MSD gene. Furthermore, we located *amhby* on the sub-telomeric region on LG24, a region showing very limited differentiation between the X and Y chromosome. The recurrent identification of *amh* duplicates as MSD genes in teleosts highlights the pivotal role of Amh signaling pathway in teleosts sex determination, and encourages further analysis on how *amh* MSDs genes initiate testicular differentiation. Moreover, our results provide an intriguing empirical example of an unexpected small sex locus with an old MSD gene and highlight the power of exciting new sequencing technologies and population genomics approaches to identify and characterize sex loci in non-model species.

## Material and Methods

### Fish rearing conditions

Research involving animal experimentation was conformed to the principles for the use and care of laboratory animals, in compliance with French and European regulations on animal welfare. The ethical agreement number for this project is 01676.02. Fertilized eggs from maturing Northern pike females were obtained from the fish production unit of the fishing federation of Ille-et-Vilaine (Pisciculture du Boulet, Feins, France). Fish were maintained indoors under strictly controlled conditions in individual aquaria to avoid cannibalism, with running dechlorinated local water (pH=8) filtered with a recirculating system. Rearing temperature and photoperiod followed the trend of the ambient natural environment. Northern pike fry were fed with live prey such as artemia larvae, daphnia, and adult artemia depending on their size. After reaching a length of 4-5 cm, Northern pike juveniles were fed with rainbow trout fry.

### Fosmid library screening, sub-cloning and assembling

The Northern pike fosmid genomic DNA library was constructed by Bio S&T (Québec, Canada) from high molecular weight DNA extracted from the liver of a male *E. lucius* from Ille et Vilaine, France, using the CopyControl Fosmid Library Production Kit with pCC1FOS vectors (Epicentre, USA) following the manufacturer’s instructions. The resulting fosmid library contained around 500,000 non-amplified clones that were arrayed in pools of 250 individual fosmids in ten 96-well plates.

Northern pike fosmid clones were screened by PCR (**Table S5**) to identify individual fosmids containing *amhby* and *amha*. To sequence the two ~ 40 kb fosmids, purified fosmid DNA was first fragmented into approximately 1.5 kb fragments using a Nebulizer kit supplied in the TOPO shotgun Subcloning kit (Invitrogen, Carlsbad, CA), and then sub-cloned into pCR4Blunt-TOPO vectors. Individual plasmid DNAs were sequenced from both ends with Sanger sequencing using M13R and M13F primers. The resulting sequences were then assembled using ChromasPro version 2.1.6 (Technelysium Pty Ltd, South Brisbane, Australia).

### Phylogenetic and synteny analyses

The protein and cDNA sequences of *amhby* and *amha* were predicted from their genomic sequences using the FGENESH+ suite [64], and the resulting cDNA sequences were compared to the corresponding transcripts publicly available in the Phylofish database [38]. Shared identity between cDNA sequences was calculated after alignment using Clustal Omega [65] implemented on EMBL-EBI [66,67] with default settings.

Global pairwise alignment of the two transcripts was performed using the mVISTA LAGAN program with default settings [68]. Shared identity and similarity between protein sequences were calculated using EMBOSS Water [69]. For each *amh* paralog, we compared the 5 kb genomic sequence upstream of the start codon using PromoterWise [69] and mVISTA LAGAN [68]. For *amha*, this 5 kb upstream region included the entire intergenic sequence up to and including part of the *dot1l* gene, located 4.3 kb from the start codon of *amha*, so the promoter for *amha* is likely located in this region.

Phylogenetic relationship reconstruction of Amh proteins was performed using the maximum likelihood method implemented in PhyML 3.0 [70], and the proper substitution model was determined with PhyML-SMS [71]. Amh protein sequences from eight teleost species and from the spotted gar (*Lepisosteus oculatus)*, which was used as outgroup, were retrieved from the NCBI Protein database. The accession number for each sequence is referenced in **Table S4**.

A synteny map of the conserved genes in blocks around *amh* was constructed with nine teleost species and spotted gar as outgroup. For spotted gar, zebrafish (*Danio rerio*), Atlantic cod (*Gadus morhua*), three-spined stickleback (*Gasterosteus aculeatus*), Japanese pufferfish (*Takifugu rubripes*), tetraodon (*Tetraodon nigroviridis*), Nile tilapia (*Oreochromis niloticus*), Southern platyfish (*Xiphophorus maculatus*) and Amazon molly (*Poecilia formosa*), the synteny map was created with the Genomicus website (<u>www.genomicus.biologie.ens.fr</u>, accessed in July 2017 [72,73]). For Northern pike, genes located upstream and downstream of *amha* on LG08 were deduced based on the location of each gene in the genome assembly (Eluc_V3, GenBank assembly accession: GCA_000721915.3).

### DNA extraction and genotyping

Small fin clips were taken from caudal fins of Northern pikes after administrating fish anesthetics (3 ml of 2-phenoxyethanol per L). When needed, fish were euthanized with a lethal dose of anesthetics (10 ml of 2-phenoxyethanol per L). All fin clips were collected and stored at 4°C in 75-100% ethanol until DNA extraction.

To obtain DNA for genotyping, fin clips were lysed with 5% Chelex and 25 mg of Proteinase K at 55°C for 2 hours, followed by incubation at 99°C for 10 minutes. After brief centrifugation, the supernatant containing genomic DNA was transferred to clean tubes without the Chelex beads [74]. Primers for genotyping were designed using Perl-Primer (version 1.1.21, [75]). Primer sequences and corresponding experiments can be found in **Table S5**.

To obtain DNA for Sanger and Illumina short read sequencing, DNA was extracted from fin clips using NucleoSpin Kits for Tissue (Macherey-Nagel, Duren, Germany) following the producer’s protocol. To obtain high molecular weight DNA for long read sequencing, DNA was extracted from blood samples lysed in TNES-Urea buffer (TNES-Urea: 4 M urea; 10 mM Tris-HCl, pH 7.5; 125 mM NaCl; 10 mM EDTA; 1% SDS) [76] followed by phenol-chloroform extraction [77]. Afterwards, the DNA pellet was washed three times with 80% ethanol before being stored at 4°C in 80% ethanol.

When preparing DNA for sequencing, DNA concentration was first quantified with a NanoDrop ND2000 spectrophotometer (Thermo scientific, Wilmington, DE) to estimate the range of concentration, and was then measured again with Qubit3 fluorometer (Invitrogen, Carlsbad, CA) to determine the final concentration.

### RNA extraction, cDNA synthesis and qPCR

Ten Northern pikes from a local French population were sampled at 54, 75, 100, and 125 days post fertilization (dpf). For the first three time points, the whole trunk, defined as the entire body without head and tail, was collected for RNA extraction because gonads were too small to be dissected routinely at these stages. At 125 dpf, gonads were large enough to be isolated and were thus collected for RNA extraction. The genotypic sex of each animal was determined based on *amhby* amplification and results are listed in **Table S6**.

Samples (trunks or gonads) were immediately frozen in liquid nitrogen and stored at - 70°C until RNA extraction. RNA was extracted using Tri-Reagent (Molecular Research Center, Cincinnati, OH) following the manufacturer’s protocol and RNA concentration was quantified with a NanoDrop ND2000 spectrophotometer (Thermo scientific, Wilmington, DE). Reverse transcription was performed by denaturing a mix of 1µg of RNA and 5 µL of 10 mM dNTP at 70°C for 6 minutes, followed by 10 minutes on ice. Random hexamers and M-MLV reverse transcriptase (Promega, Madison, WI) were then added, and the mixture was incubated at 37°C for 75 minutes followed by 15 minutes of enzyme inactivation at 70°C, and then chilled at 4°C. During these last two steps, a negative control without reverse transcriptase was prepared for each sample. The resulting cDNA was diluted 25-fold before qPCR.

Primers for qPCR were designed using Perl-Primer (version 1.1.21, [75]) on intron-exon junctions to avoid genomic DNA amplification (primers are listed in **Table S5**). Primer pairs were diluted to 6 µg/µL for qPCR, which was performed with SYBER GREEN fluorophore kits (Applied Biosystems, Foster City, CA) on an Applied Biosystems StepOne Plus instrument. For each reaction, 4 µL of diluted cDNA and 1 µL of diluted primers were added to 5 µL of SYBER Green master mix. The following qPCR temperature cycling regime was used: initial denaturation at 50°C for 2 minutes and then at 95°C for 2 minutes, followed by 40 PCR cycles at 95°C for 15s, 60°C for 30s and 72°C for 30s, and a final dissociation step at 95°C for 3s, 60°C for 30s and 95°C for 15s. Primer pairs were checked for nonspecific amplification and primer-dimers using the dissociation curve implemented in the StepOne software. Relative abundance of each target cDNA was calculated from a standard curve of serially diluted pooled cDNA from all samples, and then normalized with the geometric mean of the expression of six housekeeping genes (*β-actin, ef1α, gapdh, eftud2, ubr2*, and *ccdc6b*) following classical qPCR normalization procedures [78].

### Whole mount *in situ* hybridizations

*In situ* hybridization RNA probes for *amhby* and *amha* were synthesized from cDNA PCR products amplified from 125 dpf testis samples using primers including T7 sequences in the reverse primer (**Table S5**). These PCRs were performed with the Advantage2 Taq Polymerase (Clontech, Mountain View, CA) for high fidelity. Gel electrophoresis was performed on the PCR products, and products of the expected size were cut out and purified using NucleoSpin Gel and PCR Clean-up Kit (Macherey-Nagel, Duren, Germany). 10 ng of purified PCR product was then used as template for RNA probe synthesis. RNA probes were synthesized using T7 RNA polymerase (Promega, Harbonnieres, France) following the manufacturer’s protocol, with digoxigenin-11-UTP for *amha* and Fluorescein-12-UTP for *amhby*. Afterwards, probes were purified using NucleoSpin Gel and PCR Clean-up Kit (Macherey-Nagel, Duren, Germany), re-suspended in 50 µL of DEPC water, and stored at −70 °C.

Samples used for *in situ* hybridization were entire trunks of males and females collected at 85 dpf and fixed overnight at 4°c in 4% paraformaldehyde solution, then wash and stored at −20°c in 100% methanol. Hybridization was performed using an *in situ* Pro intavis AG robotic station according to the following procedure: male and female samples were rehydrated, permeabilized with proteinase K (25 mg/ml) for 30 minutes at room temperature, and incubated in post-fix solution (4% paraformaldehyde and 0.2% glutaraldehyde) for 20 minutes. Then, samples were incubated for 1 h at 65 °C in a hybridization buffer containing 50% formamide, 5% SCC, 0.1% Tween 20, 0.01% tRNA (0.1 mg/ml), and 0.005% heparin. RNA probes were added and samples were left to hybridize at 65 °C for 16 h. Afterwards, samples were washed three times with decreasing percentages of hybridization buffer, incubated in blocking buffer (Triton .1%, Tween 20 0.2%, 2% serum/PBS) for 2 h, and incubated for 12 h with addition of alkaline phosphatase coupled with anti-digoxigenin antibody for *amha* and anti-Fluorescein antibody for *amhby* (1:2000, Roche Diagnostics Corp, Indianapolis, IN). Samples were then washed with PBS solution, and color reactions were performed with NBT/BCIP (Roche Diagnostics Corp, Indianapolis, IN). After visual inspection of coloration, samples were dehydrated and embedded in plastic molds containing paraffin with a HistoEmbedder (TBS88, Medite, Burgdorf, Germany) instrument. Embedded samples were sectioned into 5 µm slices using a MICRO HM355 (Thermo Fisher Scientific, Walldorf, Germany) instrument. Imaging of the slides was performed with an automated microscope (Eclipse 90i, Nikon, Tokyo, Japan).

### TALENS knock-out

Three pairs of TALENs [79,80], called T1, T2 and T3, were designed to target exon 1 and exon 2 of *amhby* (**Fig S4A**). TALENs were assembled following a method derived from Huang *et al.* [81]. For each subunit, the target-specific TALE DNA binding domain consisted of 16 RVD repeats obtained from single RVD repeat plasmids kindly provided by Bo Zhang (Peking University, China). Assembled TALE repeats were subcloned into a pCS2 vector containing appropriate Δ152 Nter TALE, +63 Cter TALE, and FokI cDNA sequences with the appropriate half-TALE repeat (derived from the original pCS2 vector [81]). TALEN expression vectors were linearized by *NotI* digestion. Capped RNAs were synthesized using the mMESSAGE mMACHINE SP6 Kit (Life Technologies, Carlsbad, CA) and purified using the NucleoSpin RNA II Kit (Macherey-Nagel, Duren, Germany).

Groups of embryos at the one-cell stage were microinjected with one pair of TALENs at the concentration of either 50 ng/L or 100 ng/L. Among surviving embryos, 66 were injected with T1 (51 embryos at 50 ng/L, 15 embryos at 100 ng/L); 101 were injected with T2 (73 embryos at 50 ng/L, 28 embryos at 100 ng/L); and 45 were injected with T3 (all at 50 ng/L). At two months post fertilization, fin clips were collected from 36 surviving animals for genotyping with primers flanking the TALENs targets. Amplification of primers flanking the TALENs targets on *amhby* was also used for sex genotyping. Only genetic males with a disrupted *amhby* sequence were raised until the following reproductive season. The sperm of three one-year old G0 mosaic phenotypic males with a disrupted *amhby* sequence was collected and then used in *in vitro* fertilization with wild-type eggs. G1 individuals were genotyped for *amhby* TALEN targeting sites using primers flanking the TALENs targets, and XY mutants were kept until 153 dpf. G1 *amhby* mutants were then euthanized and dissected, and their gonads were subjected to histological analysis to inspect the phenotypes resulting from *amhby* knockout.

### Additive transgenesis with *amhby* fosmid

Northern pike embryos from wild-type parents were microinjected with the *amhby* fosmid at a concentration of 100 ng/L at the one-cell stage. Two months post fertilization, fin clips were collected and used for genotyping with primer pairs in which one primer was located on the fosmid vector sequence, which does not come from the Northern pike genome, and the other primer was located in the insert sequence from Northern pike genomic DNA to ensure specificity (**Table S5**). Only G0 males possessing the *amhby* fosmid were kept until the following reproductive season when spawning was induced with Ovaprim (Syndel, Ferndale, Washington) following the protocol from [82]. Sperm from two G0 males possessing the *amhby* fosmid was collected and used for *in vitro* fertilization with wild-type eggs. F1 XX individuals were identified due to the absence of amplification of Y-specific sequences outside of the sequence contained in the *amhby* fosmid. The XX transgenics with *amhby* fosmid were kept until 155 dpf together with wild-type sibling control genetic males and females and then dissected. Gonads from mutant and control fish were subjected to histological analysis.

### Histology

Gonads to be processed for histology were fixed immediately after dissection in Bouin’s fixative solution for 24 hours. Samples were dehydrated with a Spin Tissue Processor Microm (STP 120, Thermo Fisher Scientific, Walldorf, Germany) and embedded in plastic molds in paraffin with a HistoEmbedder (TBS88; Medite, Burgdorf, Germany). Embedded samples were cut serially into slices of 7 µm using a MICRO HM355 (Thermo Fisher Scientific, Walldorf, Germany) and stained with Hematoxylin. Imaging was performed with an automated microscope (Eclipse 90i, Nikon, Tokyo, Japan).

### Statistical analyses

All statistical analyses, including Wilcoxon signed rank test and Chi-squared test, were performed with R (version 3.5.1 [83]).

### Population analysis for male and female *E. lucius* RAD-Seq markers

A Restriction Associated DNA Sequencing (RAD-Seq) library was constructed from genomic DNA extracted from the fin clips of two parents, 37 male offspring and 41 female offspring according to standard protocols [84]. The phenotypic sex of the offspring was determined by histological analysis of the gonads at 8-month post fertilization. The library was sequenced in one lane of Illumina HiSeq 2500. Raw reads were analyzed with the Stacks (Catchen et al., 2011) program version 1.44. Quality control and demultiplexing of the 169,717,410 reads were performed using the *process_radtags.pl* script with all settings set to default. In total, 128,342,481 (76%) reads were retained after this filtering step, including ~ 1.6 M. retained sequences from the father, ~ 0.9 M. retained sequences from the mother, and between 1.0 M. and 2.2 M. retained sequences from each offspring.

Demultiplexed reads were mapped to the genome assembly of *E. lucius* (Elu_V3, GenBank assembly accession: GCA_000721915.3) using BWA (version 0.7.15-r1140, [86]) with default settings. The resulting BAM files were run through the *ref_map.pl* pipeline with default settings except a minimum stack depth of 10 (m=10). Results of *ref_map.pl* were analyzed with *populations* using the --fstats setting to obtain population genetic statistics between sexes. Fisher’s exact test was performed on all polymorphic sites using Plink (version 1.90b4.6 64-bit, [87]) to estimate association between variants and phenotypic sex. A Manhattan plot was constructed in R with homemade scripts showing -log_10_ (Fisher’s test *p*-value) for all 25 LGs of the *E. lucius* genome.

### Genome sequencing and assembly

The genome of one local phenotypic male Northern pike (*Esox lucius*) was assembled using Oxford Nanopore long reads and polished with Illumina reads.

Illumina short reads libraries were built using the Truseq nano kit (Illumina, ref. FC-121-4001) following the manufacturer’s instructions. First, 200 ng of gDNA was briefly sonicated using a Bioruptor sonication device (Diagenode, Liege, Belgium), end-repaired and size-selected on beads to retain fragments of size around 550 bp, and these fragments were A-tailed and ligated to Illumina’s adapter. Ligated DNA was then subjected to eight PCR cycles. Libraries were checked with a Fragment Analyzer (AATI) and quantified by qPCR using the Kapa Library quantification kit. Libraries were sequenced on one lane of a HiSeq2500 using the paired end 2×250 nt v2 rapid mode according to the manufacturer’s instruction. Image analysis was performed with the HiSeq Control Software and base calling with the RTA software provided by Illumina.

Nanopore long reads libraries were prepared and sequenced according to the manufacturer’s instruction (SQK-LSK108). DNA was quantified at each step using the Qubit dsDNA HS Assay Kit (Invitrogen, Carlsbad, CA). DNA purity was checked using a NanoDrop ND2000 spectrophotometer (Thermo scientific, Wilmington, DE) and size distribution and degradation were assessed using a Fragment analyzer (AATI) High Sensitivity DNA Fragment Analysis Kit. Purification steps were performed using AMPure XP beads (Beckman Coulter). In total, five flowcells were sequenced. For each flowcell, approximately 7 µg of DNA was sheared at 20 kb using the megaruptor system (Diagenode, Liege, Belgium). A DNA damage repair step was performed on 5 µg of DNA, followed by an END-repair and dA tail of double stranded DNA fragments, and adapters were then ligated to the library. Libraries were loaded on R9.4.1 flowcells and sequenced on a GridION DNA sequencer (Oxford Nanopore, Oxford, UK) at 0.1 pmol for 48 h.

In total, 1,590,787 Nanopore reads corresponding to 17,576,820,346 nucleotides were used in the assembly. Adapters were removed using Porechop (version 0.2.1, https://github.com/rrwick/Porechop). The assembly was performed with Canu (version 1.7 [88]) using standard parameters, with genomeSize set to 1.1g to match theoretical expectations [89] and maxMemory set to 240. Two rounds of polishing were performed with racon version 1.3.1 using standard parameters. For this step, Nanopore long reads were mapped to the assembly with Minimap2 (version 2.5 [90]) using the map-ont parameter preset. Afterwards, four additional rounds of polishing were performed with Pilon [91] using default parameters and the Illumina short reads mapped to the assembly with BWA mem (version 0.7.17 [86]). Metrics for the resulting assembly were calculated with the assemblathon_stats.pl script [92]. The assembly’s completeness was assessed with BUSCO [93] using the *Actinopterygii* gene set (4,584 genes) and the default gene model for Augustus. The same analysis was performed on the Elu_V3 reference genome (GenBank assembly accession: GCA_000721915.3) for comparison.

### Sequencing of male and female pools

DNA from 30 males and 30 females from the fish production unit of the fishing federation of Ille-et-Vilaine (Pisciculture du Boulet, Feins, France) was extracted with a NucleoSpin Kit for Tissue (Macherey-Nagel, Duren, Germany) following the manufacturer’s instructions. DNA concentration was quantified using a Qubit dsDNA HS Assay Kits (Invitrogen, Carlsbad, CA) and a Qubit3 fluorometer (Invitrogen, Carlsbad, CA). DNA from different samples was normalized to the same quantity before pooling for male and female libraries separately. Libraries were constructed using a Truseq nano kit (Illumina, ref. FC-121-4001) following the manufacturer’s instructions. DNAseq shorts reads sequencing was performed at the GeT-PlaGe core facility of INRA Toulouse, France (http://www.get.genotoul.fr). Two DNA poolseq libraries were prepared using the Illumina TruSeq Nano DNA HT Library Prep Kit (Illumina, San Diego, CA) following the manufacturer’s protocol. First, 200ng of DNA from each sample (male pool and female pool) was briefly sonicated using a Bioruptor sonication device (Diagenode, Liege, Belgium), and then end-repaired and size-selected on beads to retain fragments of size around 550 bp, and these fragments were A-tailed and ligated to indexes and Illumina’s adapter. Libraries were checked with a Fragment Analyzer (Advanced Analytical Technologies, Inc., Ankeny, IA) and quantified by qPCR using the Kapa Library Quantification Kit (Roche Diagnostics Corp, Indianapolis, IN). Sequencing was performed on a NovaSeq S4 lane (Illumina, San Diego, CA) using paired-end 2×150 nt mode with Illumina NovaSeq Reagent Kits following the manufacturer’s instruction. The run produced 129 millions of read pairs for the male pool library and 136 millions of read pairs for the female pool library.

### Identification of sex-differentiated regions in the reference genome

Reads from the male and female pools were aligned separately to the reference genome (GenBank assembly accession: GCA_000721915.3) using BWA mem version 0.7.17 [86] with default parameters. Each resulting BAM file was sorted and PCR duplicates were removed using Picard tools version 2.18.2 (http://broadinstitute.github.io/picard) with default parameters. Then, a pileup file combining both BAM files was created using samtools mpileup version 1.8 [94] with per-base alignment quality disabled (-B). A sync file containing the nucleotide composition in each pool for each position in the reference was generated from the pileup file using popoolation mpileup2sync version 1.201 [95] with a min quality of 20 (--min-qual 20).

An in-house C++ software was developed to identify sex-specific SNPs, defined as positions heterozygous in one sex while homozygous in the other sex, compute per base between-sex F_ST_, and coverage for each sex from the output of popoolation mpileup2sync (PSASS v1.0.0, doi: 10.5281/zenodo.2538594). PSASS outputs 1) all positions with sex-specific SNPs or high F_ST_, 2) the number of such positions in a sliding window over the genome, 3) average absolute and relative coverage for each sex in a sliding window over the genome, and 4) number of sex-specific SNPs as well as coverage for each sex for all genes and CDS in a user-supplied GFF file.

We used PSASS to identify non-overlapping 50 kb windows enriched in sex-specific SNPs in *E. lucius*, using the following parameters: minimum depth 10 (--min-depth 10), allele frequency for a heterozygous locus 0.5 ± 0.1 (--freq-het 0.5, --range-het 0.1), allele frequency for a homozygous locus 1 (--freq-het 1, --range-het 0), --window-size 50000, and --output-resolution 50000. For scaffold1067, the number of sex-specific SNPs and coverage for each sex were similarly computed in 2,500 bp windows, only changing the parameters --window-size to 2500 and --output-resolution to 2500.

### Identification of Y specific sequences in the Nanopore assembly

A sync file containing the nucleotide composition in each pool for each position in the reference was generated as described in the previous section, using the Nanopore assembly (NCBI accession number SDAW00000000) to align the reads. This time, because we were comparing reads coverage levels, the BAM files were filtered with samtools version 1.8 [94] to only retain reads with a properly mapped pair and a mapping quality higher than 30 to reduce the impact of false positive mapped reads. The resulting sync file was used as input for PSASS to compute coverage for each sex in 1 kb non-overlapping windows along the genome using the following parameters: --min-depth 10, --window-size 1000, and --output-resolution 1000. Results from this analysis were filtered in R (version 3.5.1 [83]) to identify 1 kb regions with mean relative coverage between 0.3 and 0.7 in males, and mean absolute coverage lower than 1 in females.

To identify protein coding sequences, we performed alignments between the Y-specific sequence and the teleostei (taxid:32443) non-redundant protein database using blastx (https://blast.ncbi.nlm.nih.gov/Blast.cgi, version 2.8.1 [96]) with the parameters “Max target sequences” set to 50 and “Max matches in a query range” set to 1. Regions matching potential homologs with an e-value < 1E-50 were considered as protein coding sequences.

To determine repeat content of the Y-specific sequence, RepeatMasker (version open 4.0.3 [97]) was run on these Y-specific sequences with NCBI/RMBLAST (version 2.2.27+) against the Master RepeatMasker Database (Complete Database: 20130422). The same analysis was also performed on the entire Nanopore assembly.

## Supporting information

Supplementary file 1, Figure and Text: Figure S1 to Figure S6 and additional results. Supplementary file 2, Tables: Table S1 to Table S8

## Data availability

Sequencing data and assembly for the *Esox lucius* genome can be found under NCBI Bioproject PRJNA514887. Sequencing data for identifying and charactering the sex locus, including RAD-seq, pool-seq reads, can be found under the NCBI Bioproject PRJNA514888.

### Acknowledgement

We are grateful to the fish facility of INRA LPGP for support in experimental installation and fish maintenance. We are grateful to the genotoul bioinformatics platform Toulouse Midi-Pyrenees (Bioinfo Genotoul) for providing help and/or computing and/or storage resources. We also want to thank Nicolas Perrin, Susana Coelho and Eric Pailhoux for feedbacks and helpful discussion on an earlier version of the manuscript. This project was supported by funds from the Agence nationale de la recherché (France) to Y.G. and the Deutsche Forschungsgemeinschaft to M.S. (ANR/DGF, PhyloSex project 2014-216). The GeT, Toulouse (CR, JL), and the MGX France (LJ, HP) core facilities were supported by France Génomique National infrastructure, funded as part of “Investissement d’avenir” program managed by Agence Nationale pour la Recherche (contract ANR-10-INBS-09). The funders had no role in study design, data collection and analysis, decision to publish, or preparation of the manuscript.

## Author contribution

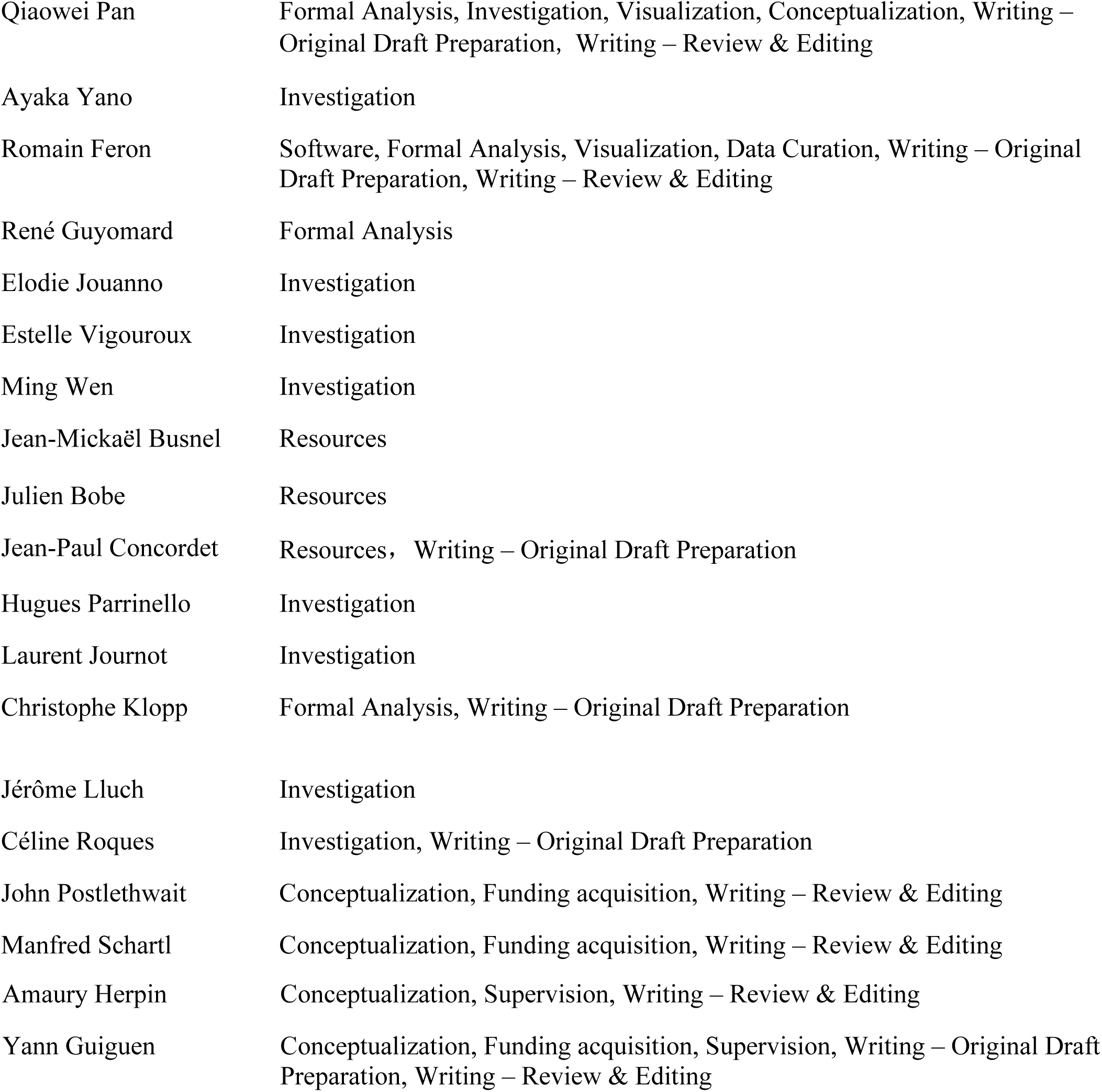

